# Metabarcoding for parallel identification of species, sex and diet: an application to the conservation of globally-threatened *Gyps* vultures

**DOI:** 10.1101/756247

**Authors:** Mousumi Ghosh-Harihar, Nehal Gurung, Harsh Shukla, Ishani Sinha, Awadhesh Pandit, Vibhu Prakash, Rhys E. Green, Uma Ramakrishnan

## Abstract

An understanding of the factors affecting the diet composition of critically endangered *Gyps* vultures in the Indian subcontinent has important applications to the design of effective conservation strategies. After suffering a massive decline (~99%) in numbers caused by veterinary use of the nephrotoxic drug diclofenac, vultures now persist in very low numbers, mostly concentrated within or near National Parks. This spatial association might be attributed to the availability of wild ungulate carcasses free from toxic veterinary drugs. Hence, quantification of vulture diets and the factors affecting them is critical to test this hypothesis. Here, we describe and validate a robust “field-to-benchtop-to-desktop” metabarcoding workflow for assessing the species- and sex-specific diet of these obligate scavengers from non-invasively collected faecal samples. Seven metabarcodes targeting mitochondrial and nuclear genes were used to simultaneously determine the vulture species identity, sex and species consumed. The amplicons were sequenced using an Illumina Miseq platform. We included controls and three replicates per sample to establish a series of non-arbitrary thresholds to filter the sequence data and eliminate cross-contamination, PCR/sequencing errors and false positives. Using this strategy enabled identification of species and sex for all samples. Diet species-specific sequences could be identified with high taxonomic resolution for 93% of samples. Out of the seven metabarcodes, just four were sufficient to meet the objectives. From this preliminary analysis, domestic livestock seem to be the most frequently consumed diet item across samples from inside and outside protected habitats. Our method provides a rapid and reliable tool for describing large-scale variation in consumption of domestic versus wild species in the diet of these species, paving the way for a better understanding of the role protected areas play in persistence and recovery of the remaining *Gyps* vulture populations in the wild.

## 1. Introduction

DNA metabarcoding coupled with high-throughput-sequencing (HTS) has revolutionized our ability to investigate the dietary ecology of species (Pompanon et al., 2012; Taberlet et al., 2012). The cost-effectiveness and relative ease of implementation has made diet metabarcoding an increasingly popular approach (Soininen et al., 2015; Zinger et al., 2019). Compared to morphological or micro-histological analyses of food remains or traditional molecular approaches, metabarcoding allows processing a large number of samples quickly and provides greater taxonomic coverage and resolution in a single run (Deagle et al., 2014; Tillmar et al., 2013; Wirta et al., 2014). However, obtaining robust inferences from metabarcode data is challenging because multiple biases may occur anytime from fieldwork to bioinformatic analyses. Such biases could result in false negatives, false positives and low taxonomic resolution (Ficetola et al., 2016; Galan et al., 2018; Zinger et al., 2019).

For instance, contamination during sample collection, extraction and laboratory processing may be amplified further due to PCR and high-depth of sequencing in HTS. Incorrect design of primers or choice of metabarcode region could lead to no or biased/differential amplification of taxa, or result in poor taxonomic resolution (Clarke et al., 2014; De Barba et al., 2014; Deagle et al., 2014; Pompanon et al., 2012). Since DNA derived from food items is often degraded in faecal samples, the use of relatively short mitochondrial markers and multiplexing regions can be used to minimize false negatives (Deagle et al., 2006). Similarly, primer bias could be a major source of false negatives. False positives can be introduced at various stages due to reagent contamination or cross contamination during sample collection, extraction or PCR (Taberlet et al., 2018). Another source of false positives is ‘index-jumping’ (involving Illumina sequencing adapter when several libraries are pooled together) or ‘mistagging’ (tags recombining with unintended sample) (Schnell et al., 2015). This form of sample cross-contamination usually involves the most abundant taxa, but can skew the results for samples with low DNA concentrations (Esling et al., 2015; Schnell et al., 2015). Furthermore, a limitation of metabarcoding compared to traditional diet analysis remains the inability to convert sequence data into relative biomass of different diet items consumed owing to PCR biases (Elbretch & Leese 2015).

Several approaches have been suggested in recent reviews to minimize these biases to allow sound ecological inferences from metabarcoding studies (Zinger et al., 2019). In general, the emphasis is on including several types of experimental controls to identify and exclude noise from the HTS data generated. The suggested experimental controls include: (a) pilot experiments to assess sampling design (Dickie et al., 2018); (b) sufficient biological and technical replicates (Ficetola et al., 2015); (c) negative controls at various experimental stages (sampling, extraction, PCR, and sequencing) and positive controls (mock DNA communities; e.g. (Corse et al., 2017)) for data validation. All these controls help detect contamination, identify filtering and clustering thresholds, thereby ensuring reliability of the entire workflow (De Barba et al., 2014). Finally, during bioinformatic analyses, adopting filtering steps relevant to experimental design and ecological question is critical for making robust inferences (Ficetola, Taberlet, and Coissac 2016; Galan et al. 2016). For instance, non-arbitrary experimental controls could be used to ascertain filtering thresholds, providing a direct measurement of artefacts derived from PCR/sequencing/tagging errors (Corse et al., 2017; Galan et al., 2018). Additionally, taxonomic assignment should incorporate a priori ecological knowledge of the plausible diet taxa in the ecosystem where sampling happens. This could be further augmented by a robust reference database of metabarcode sequences of expected species when possible (e.g. Alsos et al. 2018).

Generating reliable dietary information is a particularly critical requirement for conservation of three species of *Gyps* vultures (White-rumped vulture *Gyps bengalensis*, Long-billed vulture *G. indicus* and Slender-billed vulture *G. tenuirostris*) in the Indian subcontinent. These species were listed by IUCN as Critically Endangered following a catastrophic decline in their numbers caused by veterinary use of the nephrotoxic non-steroidal anti-inflammatory drug (NSAID) diclofenac (Green et al., 2004; Oaks et al., 2004; Prakash et al., 2003). While population monitoring indicates that populations are stabilizing and even recovering in the aftermath of the ban on the veterinary use of the drug, most remaining vultures are found inside or near National Parks (Chaudhary et al., 2012; Prakash et al., 2012, 2019; Galligan et al. 2019). It has been hypothesized that high levels of consumption of food from carcasses of wild ungulates, which are not contaminated with NSAIDs, in such areas might be responsible for the higher numbers of vultures observed in and near National Parks. But testing this assumption requires robust data on diets of wild vultures. This is especially the case because some protected areas, including National Parks, do not support large populations of wild ungulates. A better understanding of spatial variation in the relative contribution of ‘safe’ food from wild ungulates in vulture diet will therefore aid in identifying areas where conservation efforts, including reintroduction of captive-bred vultures can be prioritized. This is particularly important since illegal use of diclofenac continues, alongside legal use of several other drugs now established to be nephrotoxic to vultures (Cuthbert et al., 2016).

In the past, vulture diet has been studied using morphological and micro-histological examination of undigested food remains (hair, bones) from regurgitated pellets (Donázar et al., 2010; Kelly et al., 2007). This approach is likely to be biased in the Indian subcontinent, given the frequent practice of skinning domesticated ungulate carcasses before disposal. Furthermore, because our long-term goal is to study differences in the diet of vultures over large areas, assessing diet by using non-invasive methods such as the collection of faecal droppings is the most practical approach. A previous study examining the gut microbiome of New World Vultures using metabarcoding also included a single mammalian primer set (16Smam1 and 16Smam2) to detect diet DNA (Roggenbuck et al., 2014). They found that only 8% of the hindgut samples yielded any mammalian DNA. This was attributed to the extremely harsh chemical conditions in the gastrointestinal tract, because DNA from food items could be detected from facial swabs in 90% of the samples. Therefore, establishing a robust protocol for dietary metabarcoding using suitable markers, which can recover diet metabarcodes even from highly degraded faecal samples, is of critical importance.

In this study, we describe a rigorous “fieldwork-to-benchtop-to-desktop” metabarcoding workflow to determine the species- and sex-specific diet of *Gyps* vultures from their faeces. We had two specific aims. Firstly, to establish data reliability in order to minimize false-positives and false-negatives. Towards this end, we used a robust experimental design including positive and negative controls as well as technical replicates to identify clustering-free and non-arbitrary thresholds (based on controls) to filter sequence occurrences (Corse et al., 2017; Ficetola et al., 2016; Galan et al., 2018). Secondly, by applying this protocol to wild collected faecal samples, we compared seven markers in terms of identification success, taxonomic resolution and amplification biases to identify the most optimal set for simultaneously assessing the vulture species identity, sex and diet.

## 2. Materials and Methods

### 2.1 Faecal sample collection

We collected faecal samples from captive individuals held at the vulture conservation breeding centre at Pinjore, Haryana, (March 2018) and from wild vultures at sites located in Pench, Kanha, Sariska, Ranthambore and Mukundra Hills Tiger Reserves in India (June 2018-December 2018). At each field site, we located resident *Gyps* vulture (White-rumped and Long-billed) roosting and nesting sites and collected faecal droppings from substrates (soil, leaves, rocks) directly under the roost/nest. In order to minimize chances of sampling the same individual repeatedly, we collected droppings which were spatially separated. We attempted to collect relatively fresh samples (as determined visually by extent of desiccation) where possible to avoid false negatives (see Oehm et al. 2011). We swabbed the faecal dropping using a plastic applicatory rayon swab dipped in Longmire lysis buffer (Longmire et al., 1997). The tip of the swab was broken and stored in a 2 ml vial (with 1 ml buffer) at room temperature for up to 30 days, after which it was transported to the laboratory and stored at −20°C until DNA extraction.

### 2.2 Faecal DNA extraction

All faecal DNA extraction steps were carried out in a room dedicated to extracting DNA from non-invasive/degraded sources while following standard safety and contamination control precautions. DNA from faecal samples was extracted following the manufacturer’s recommendations using the QIAmp Fast DNA Stool Mini Kit (QIAGEN, Germany). Extractions were carried out in batches of 24 including a negative control to monitor potential contamination.

### 2.3 Metabarcoding primers

For parallel identification of vulture species, sex and diet, we designed specific primers targeting variable DNA regions for multiplex primer reactions using Primer 3.0 v.0.4.0 (http://bioinfo.ut.ee/primer3-0.4.0/), ensuring that the amplicon lengths (including primer sequences) were within the range of 130-150 bp (Table 1). For distinguishing between the *Gyps* species, published markers rely on amplification and restriction digestion of larger fragment of COI (806 bp) (Kapetanakos et al., 2014), which might not be usable for degraded DNA found in faecal samples. Hence, we designed two primer pairs (*CytbV1* and *CytbV2*) that amplify short fragments from the 5’ end of cytochrome b gene of the mitochondrial genome. While the use of primers targeting the chromohelicase-DNA binding gene has previously been described to sex *Gyps* vultures (Ghorpade et al., 2012), the W-specific primer targeted a longer fragment (258bp) and we therefore redesigned a set of primers targeting the ZW-common (*VsexZW*) and W-specific (*VsexW*, female) regions.

**Table 1.**
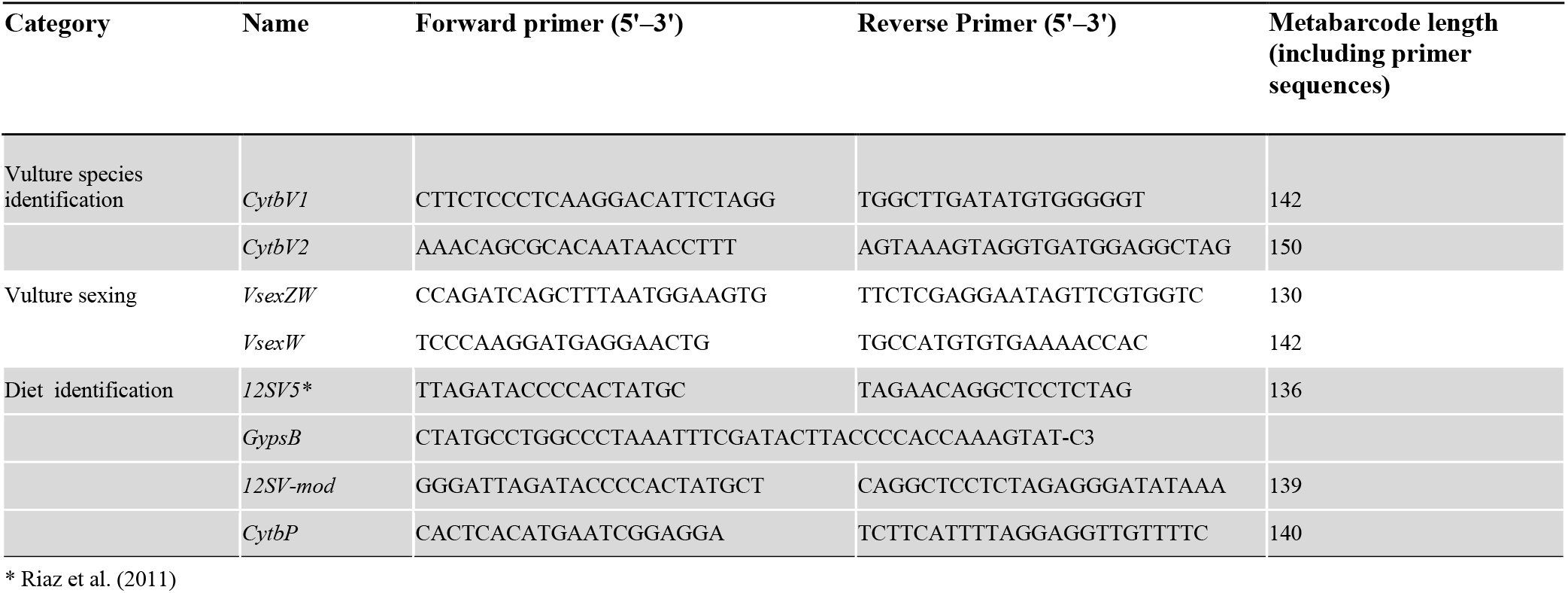
Sequences of the metabarcoding primers used in this study. The *GypsB* blocking oligo overlaps with the 12SV5 forward primer and has a C3 spacer modification at the 3’ end.

We used the primer pair *12SV5*, which targets all vertebrates, described in (Riaz et al., 2011) in order to identify all the diet species (Table S1, Figure S1). During an initial run of sequencing for six samples, we recovered barely 0.09% of diet sequences using *12SV5*, while the remaining sequences were derived from vultures. Therefore, in order to limit the amplification of *Gyps* DNA when using the *12SV5* primer, we designed a vulture-specific blocking oligonucleotide (*GypsB*), as suggested by Vestheim & Jarman (2008) (Table 1). We also used two additional primer pairs, which should not amplify *Gyps* DNA owing to mismatches with the reverse primer sequence. *12SVmod* targeted an overlapping region of the V-loop of the 12S mitochondrial gene while *CytbP* targeted the cytochrome b gene (Table 1). All primers were modified by adding the Illumina overhang adapters (forward overhang: 5’ TCGTCGGCAGCGTCAGATGTGTATAAGAGACAG-primer sequence; reverse overhang: 5’ GTCTCGTGGGCTCGGAGATGTGTATAAGAGACAG-primer sequence).

### 2.4 Positive and negative controls

An increasing number of studies emphasise the importance of including an adequate number of both positive and negative controls at the extraction, PCR amplification and sequencing steps to reduce the number of false negatives and positives (Corse et al., 2017; De Barba et al., 2014; Galan et al., 2018; Zinger et al., 2019). Hence, in our protocol, we similarly amplified two positive controls (*T_pos_*). The first one (sample-H2) consisted of a faecal sample from a captive Himalayan Griffon (*G. himalayensis*), which had been fed exclusively on goats. The second one (sample-MC1) comprised of a mix of DNA extracted from a shed feather of a Long-billed vulture (0.8 nM/ul) and equal concentration of DNA (0.2 nM/ul, determined using a Qubit fluorometer) from three putative diet species (Domestic Cow *Bos taurus*, Blackbuck *Antilope cervicapra* and Barking Deer *Muntjak muntiacus*) extracted from reference samples available in the laboratory. In addition, we included multiple negative controls (i) two extraction controls (*T_ext_*) to detect cross-sample or exogeneous contamination during DNA extraction, (ii) one PCR control per multiplex reaction (*T_pcr_*) to indicate cross-contamination during preparation of PCR mix (primers but no DNA template in the vial), and (ii) an indexing control during library preparation (*T_tag_*) to detect mis-tagging due to recombination of sequences from different samples (an empty vial with unique indexes, but no primers or DNA template) (Corse et al., 2017; Esling et al., 2015; Schnell et al., 2015).

### 2.5 PCR and library preparation

We amplified each of 28 wild-collected and 2 positive control samples using three different multiplex PCR reactions to avoid cross-amplification between primer pairs (Figure S2). The first multiplex set included *CytbV1, 12SV5, GypsB* and *CytbP (M1)*, the second set included *CytbV2* and *12SVmod* (*M2*) and the final set included *VsexW* and *VsexZW* primers (*M3*). We included three PCR replicates for each sample per multiplex reaction, resulting in a total of nine PCR reactions per sample. Each PCR reaction was performed in 25 μl reaction volume consisting of 12.5 μl of 2x Qiagen Multiplex Kit Master Mix (Qiagen), 1-1.25 μl of forward and reverse primers (0.12/0.14 μM of mitochondrial primers, 2 μM of blocking oligonucelotide and 0.27 μM of nuclear primers) and 2.5-5 μl of template DNA. The PCR conditions included an initial denaturation step at 95°C for 15 min, followed by 40 cycles of denaturation at 95°C for 30 sec, annealing at 60°C for 30 sec, and extension at 72°C for 20 sec, followed by a final extension step at 72°C for 5 min.

Amplification was checked using gel electrophoresis on 2% agarose gels and product concentrations were quantified using a Qubit Fluorometer (Invitrogen, Darmstadt, Germany). Then the PCR products for each sample were pooled across the three multiplex reactions (7.5 μl from *M1*, 7.5 μl from *M2* and 10 μl from *M3*), so that finally, each sample had three replicates. The pooled amplicons were purified to remove free primers and primer dimers using Agencourt AMPure XP Beads (Beckman Coulter, Fullerton, CA, USA) using 37.5 μl (1.5x) of resuspended beads and 25 μl of DNA. In the next step, we attached dual indices (Nextera XT Indexing Kit) through a limited-cycle PCR. The PCR reaction was carried out in 50ul reaction volume, using 25 μl of Qiagen Multiplex Kit Master Mix (Qiagen) and 5 μl of each indexed primer and 20ng of the purified DNA template. The PCR conditions included an initial denaturation step at 95°C for 3 min, followed by 8 cycles of denaturation at 95°C for 30 sec, annealing at 55°C for 30 sec, and extension at 72°C for 30 sec, followed by a final extension step at 72°C for 5 min. By using 12 index i5 and eight index i7, we were able to multiplex 96 samples in the same Miseq run. The PCR products were again purified using Agencourt AMPure XP Beads (Beckman Coulter, Fullerton, CA, USA) using 50 μl (1.5x) of resuspended beads and 50 μl of DNA. To verify the final library fragment size (~263bp) and detect potential primer contaminations, we analysed 1 μl (1:50 dilution) of the dual-indexed library on a 2100 Bioanalyzer and Agilent High Sensitivity DNA chip (Agilent Technologies, Santa Clara, USA).

### 2.6 Miseq Sequencing

We chose the Miseq platform, since it is known to generate lower error rates than other HTS platforms (D’Amore et al. 2016). The dual-indexed library was pooled by volume (5 nM per sample). We denatured the pooled library using NaOH, diluted it with the hybridization buffer and loaded 6 pM and 10% PhiX control on a Miseq flow cell with a 500-cycle Reagent Kit v2 (Illumina). We carried out a run of 2 x 150 bp paired-end sequencing, which yielded high-quality sequencing with each nucleotide of the seven metabarcodes read twice after assembly of read 1 and 2.

### 2.7 Sequence analyses and filtering

We used OBITools (Boyer et al., 2016) to analyse the HTS data, separately for each of the seven metabarcodes (Table 1). Using the *illuminapairedend* script, we assembled the forward and reverse reads into a single sequence. Thereafter, we demultiplexed and filtered the primers and tags using the *ngsfilter* script as described in Shehzad et al. (2012). We excluded all sequences shorter than 80 bp, or sequences with counts less than 10. PCR errors can generate variants with very high frequency when compared to error-free sequences. Hence, we used the *obiclean* script to assign each sequence to the status of “head”, “internal” or “singleton” according to a directed acyclic graph (De Barba et al., 2014). All sequence variants classified as “internal” based on a 5% threshold (determined from known variants in the mock sample) were discarded to further filter out variants resulting from PCR/sequencing errors (Table 2).

**Table 2.** Summary of the number of sequences retained after different steps of the HTS filtering protocol. Percentage of sequences retained from the previous step are indicated in parentheses.

|  | Total | Metabarcoding primers |  |  |  |  |  |  |
| --- | --- | --- | --- | --- | --- | --- | --- | --- |
|  |  | *CytbV1* | *CytbV2* | *VsexW* | *VsexZW* | *12SV5* | *12SVmod* | *CytbP* |
| Estimated paired-end sequence reads | 11,182,223 |  |  |  |  |  |  |  |
| Sequence reads for which primers were identified |  | 2,024,317 (18.1) | 1,963,151 (17.6) | 50,682 (0.5) | 3,670,957 (32.8) | 328,083 (2.9) | 246,415 (2.2) | 29,911 (0.3) |
| Unique sequences |  | 24,333 (1.2) | 21,026 (1.1) | 1,968 (3.9) | 42,346 (1.2) | 10,531 (3.2) | 5,958 (2.4) | 965 (3.2) |
| Unique sequences longer than 80bp and count >10 |  | 1,165 (4.8) | 1,431 (6.8) | 155 (7.9) | 3,471(8.2) | 967 (9.2) | 666 (11.2) | 123 (12.7) |
| Unique sequences after removal of noise/head |  | 265 (22.7) | 368 (25.7) | 28 (18.1) | 1051 (30.3) | 203 (21.0) | 136 (20.4) | 8 (6.5) |
| Unique sequences with >95% identity |  | 254 (95.8) | 363 (98.6) | 27 (96.4) | 1047 (99.6) | 190 (93.6) | 130 (95.6) | 7 (87.5) |
| Unique sequences retained after *LFNpos* |  | 18 (7.1) | 23 (6.3) | 22 (81.2) | 28 (2.7) | 162 (85.3) | 118 (90.1) | 7 (100) |
| Unique sequences retained after *LFNneg* |  | 11 (61.1) | 12 (52.2) | 2 (9.1) | 18 (64.3) | 162 (100) | 106 (89.8) | 5 (71.4) |
| Variants which were present in at least two out of three replicates |  | 6 (54.5) | 6 (50) | 2 (100) | 8 (44.4) | 125 (77.2) | 62 (58.5) | 5 (100) |
| MOTUs |  | 5 (83.3) | 3 (50) | 2 (100) | 2 (25) | 12 (9.6) | 10 (16.1) | 3 (60) |

Reference databases corresponding to each metabarcode were assembled using the program *ecoPCR* (Ficetola et al., 2010) based on sequences derived from the European Molecular Biology Laboratory (EMBL) database. Each sequence was assigned to a unique taxon using the script *ecotag*, corresponding to the last common ancestor node in the NCBI taxonomic tree of all the taxids of the sequences of the reference database that matched the query sequence. The automatically assigned taxonomic identification underwent further filtering based on multiple low-frequency noise (*LFN*) thresholds to eliminate false-positives arising likely due to contamination, mistagging and PCR/sequencing errors (Corse et al., 2017; De Barba et al., 2014). Firstly, we retained only sequences with specified identity threshold of ≥95% over the entire query sequence length with the corresponding reference sequence to increase the accuracy of the automatic taxonomic assignment and remove chimeras.

#### 1. Filtering using positive and negative controls

We then applied an *LFNpos* threshold, based on the relative frequency of the least frequent expected variant across the known and mock community replicates, and removed all variants with counts below this threshold (Corse et al., 2017). We did not get unexpected sequences across the replicates in our positive controls. To further remove low frequency noise in replicates with low numbers of reads, we used *LFN_neg_* threshold, determined for each variant by the maximum value across the negative control replicates (*T_neg_*). Read counts lower or equal to this threshold could not be distinguished from noise and were hence, replaced with zero, while we retained the read counts for the positive samples (Galan et al., 2018).

#### Filtering using replicates

Thereafter, we assessed the repeatability of the experimental procedure in two ways: by estimating the distance between replicates as suggested by De Barba et al. (2014) and by Corse et al. (2017). First, we retained only those sequence variants which were present in at least two replicates of a sample. Second, we calculated the Renkonen distance (RD) between replicates of the same sample and discarded the replicates for which the distance was more than a threshold set by 10% upper tail of the distribution of RDs. Finally, all samples with only one replicate were removed and for the rest, the read counts were added across replicates for each sample variant. However, we do not make any inference regarding relative abundance of diet species based on read counts due to possible PCR bias (Elbrecht and Leese, 2017).

### 2.8 Taxonomic assignment

We assigned the final set of variants per metabarcode to unique molecular taxonomic units (MOTU) using multiple approaches. Firstly, we collapsed variants into discrete taxa using a 2% sequence divergence threshold and relative abundance of the sequences. For sequences with >2% divergence, we performed additional phylogenetic analyses, where the topology of the trees was used to resolve the taxonomy of the variants. We also used additional BLAST hits in GENBANK with very stringent criteria (E-value threshold: 1e-10; minimum query coverage: 98%) for confirming species-level identity. Finally, the assignment was refined with biogeographic information on known diet species in the region. We also computed the identification resolution index (IR, (Corse et al., 2017; Zarzoso-Lacoste et al., 2016) for each diet metabarcode and compared them to assess their performance in detecting diet items. For this, we attributed a score to each sequence variant in every sample based on its level of taxonomic assignment (species-6, genus-5, subfamily-4, family-3, suborder/infraorder-2), with the IR for a primer representing the mean across samples. We tested for difference between diet metabarcodes using Wilcoxon rank test.

## 3. Results

### 3.1 HTS data filtering

The Miseq sequencing of the amplicon library (96 products including six negative controls and three replicates each of 28 samples collected from the wild and two positive controls) generated a total of 11,182,223 paired-end reads of the seven metabarcodes. The total sequence variants retained after each stage of filtering are reported in Table 2 for every metabarcode. In general, despite discarding a large number of sequence variants (92-99% with species identification and sexing primers; 28-52% with diet primers) the three *LFN* filters retained between 88.7-99.9% of the total reads. The maximum number of sequence reads were assigned to *VsexZW* (32.8%), while the minimum number was assigned to *CytbP* (0.3%). In general, the reads assigned to *Gyps* species identification and sexing primers were much higher (69%) than the diet-identification primers (5%) (Table 2).

### 3.2 Species identification and sexing

Between the two species identification primers, *CytbV1* performed better in terms of providing taxonomic resolution. It could identify five different species (four *Gyps* species including Himalayan vulture from captivity, Eurasian Griffon (*G. fulvus*) *and* Red-headed Vulture (*Sarcogyps calvus*)), while *CytbV2* could only distinguish White-rumped Vulture and grouped the rest into the genus *Gyps* (Fig. 2). It could assign the sample belonging to Red-headed Vulture to family Accipitridae. None of the two primers produced any sequences for sample GRM1, which was identified as Egyptian vulture *Neophron percnopterus* (belonging to the subfamily Gypaetinae unlike the others which belong to the subfamily Aegypiinae) by *12SV5*. One sample from Kanha Tiger Reserve (KNH3) contained sequences specific to both White-rumped (read count-39010) and Long-billed (read count-17972) suggesting mixing of samples. The female-specific *VsexW* amplified across 16 samples. The *VsexZW* produced sequences which could be assigned to both Z and W chromosome, separately (Fig. 2). The Z-specific sequences were retrieved for all 30 samples and the W-specific sequences using the same primer was retrieved for the same 16 samples (as with *VsexW*) and an additional one (GRM1, Egyptian vulture).

**Figure 1.**
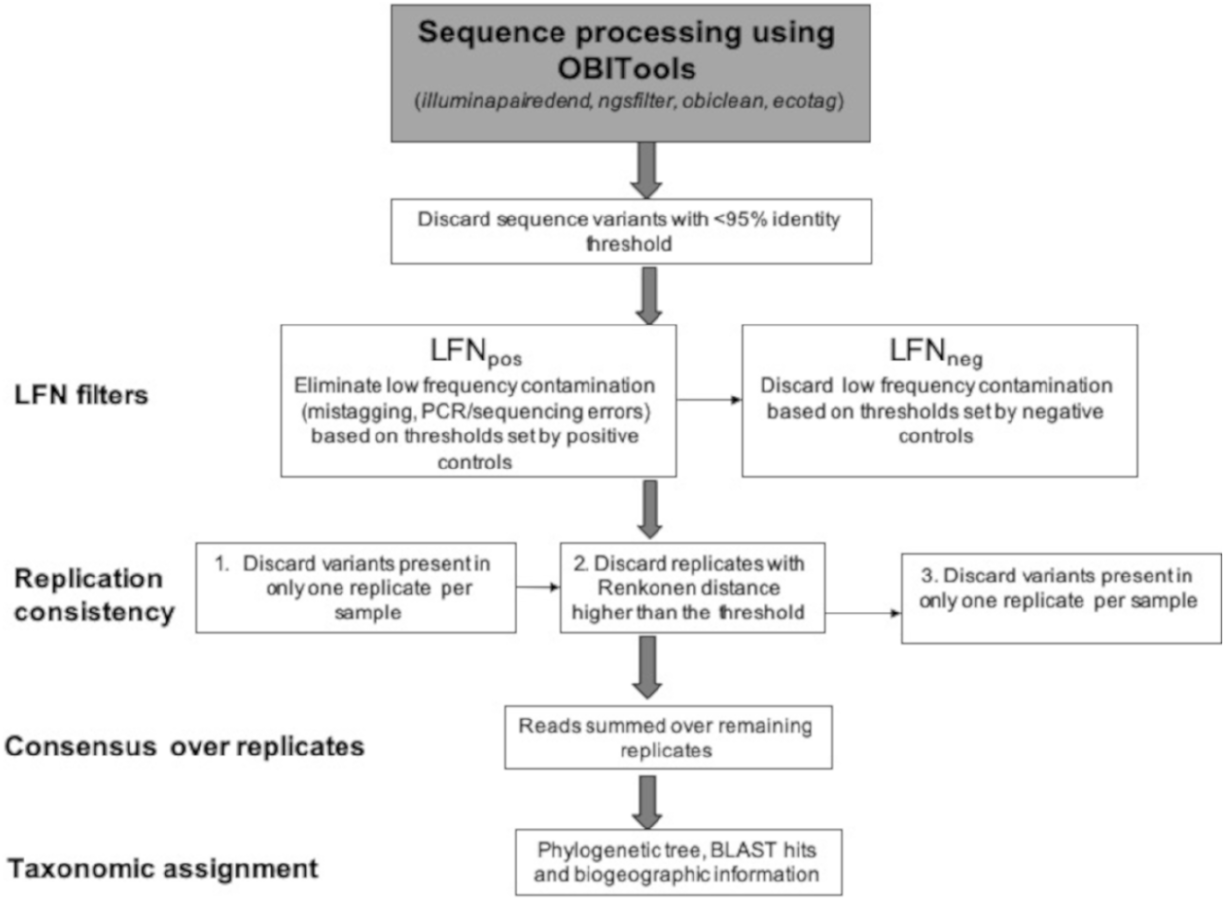
Schematic representation of the HTS filtering pipeline followed for the metabarcoding sequences.

**Figure 2.**
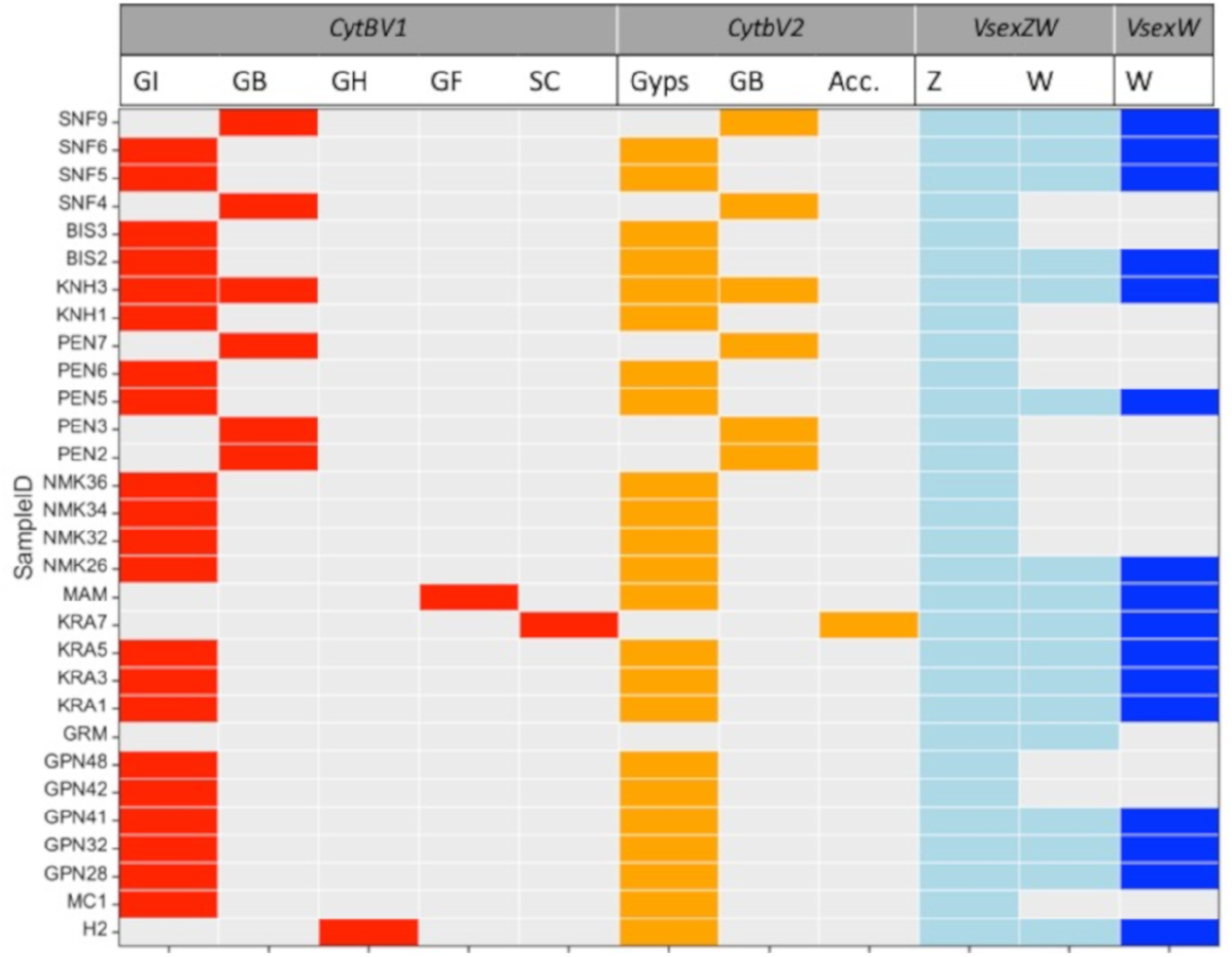
The primer-wise taxonomic and sex assignment of the samples. Abbreviations: *G-Gyps, GI-Gyps indicus*, GB-G. *bengalensis*, GH-G. *himalayensis*, GF-G. *fulvus, SC-Sarcogyps calvus*, ACC-Accipitridae, Z-Z chromosome, W-W chromosome (female-specific). One sample (KNH3) contained sequences specific to both *G. indicus* and *G. bengalensis*, making the origin of the W-specific sequences uncertain.

### 3.3 Diet species detection

Despite using a blocking oligonucleotide (*GypsB*) with *12SV5*, the amplification of vulture sequences (13.6% of total sequence reads) was completely blocked in only three samples. In half the samples the percentage of vulture sequences was less than 3%, while it was above 85% for four samples (Fig. 3a). However, sufficient diet-specific sequence reads remained in 26 samples (after HTS filtering) to allow taxonomic assignment. The modified *12SVmod* primer did not produce any vulture-specific sequences and amplified diet sequences in 28 out of 30 samples. However, as seen in similar experiments (Deagle et al., 2009; Shehzad et al., 2012), we detected some human contamination (0.06%) in the sequences corresponding to the *12SVmod* primer. Only nine samples were amplified by *CytbP*.

**Figure 3.**
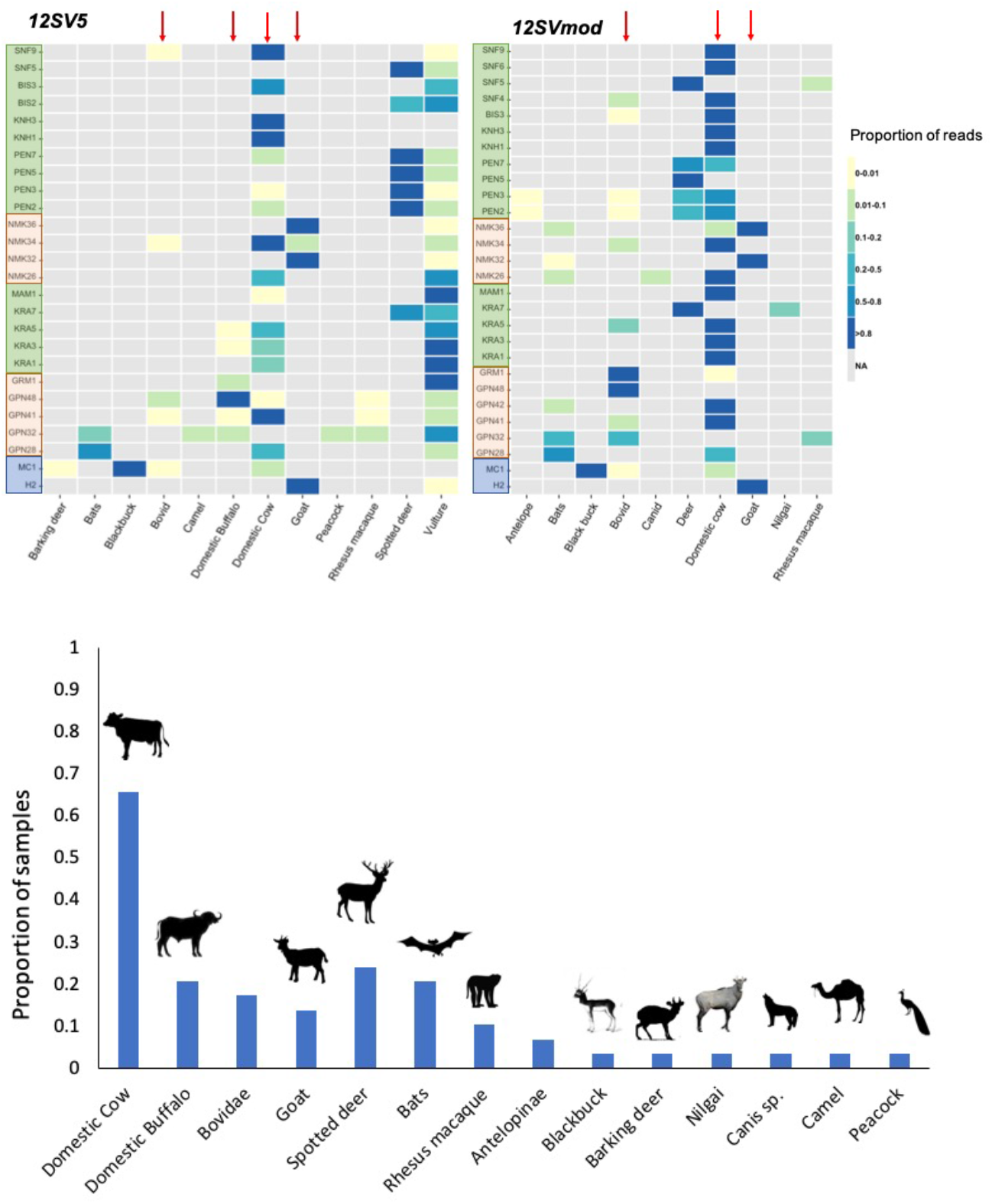
The proportion of reads assigned to each diet category across all the samples (top). The diet composition of *Gyps* vultures (bottom). The samples (on Y-axis) from inside national parks are shaded in green, those outside in orange and the positive controls are shaded in blue.

In terms of taxonomic resolution, *12SV5* performed slightly better than *12SVmod* in terms of the taxonomic IR (Wilcoxon rank test: W=553, p = 0009; Fig. 4). Firstly, it identified all three diet species in the mock community (Black buck, Domestic cow and Barking Deer), while *12SVmod* did not retrieve any sequences corresponding to barking deer or the family Cervidae (Fig. 3a). At least three samples (GPN42, SNF4, SNF6) yielded diet-specific sequences only while using *12SVmod*, and only *12SV5* identified diet remains in one sample (BIS2). When used together, they identified diet remains in 29/30 samples. One sample (PEN6) did not produce any diet-specific sequences although the vulture species and sex could be determined from the sample. In 38% of the cases, both primers identified the same taxa (Fig 3a). Three diet species were picked up by *12SV5* alone (Camel *Camelus dromedarius*, Buffalo *Bubalus bubalis*, Peacock *Pavo cristatus*), and *12SVmod* uniquely identified three taxa (Nilgai *Boselaphus tragocamelus, Canis* sp. and an Antelopinae). In seven samples, *12SV5* identified additional species, while *12SVmod* did the same in six cases (Fig. 3a). *CytbP* only identified three genera (*Axis sp., Antilope sp*. and *Capra sp*.).

**Figure 4.**
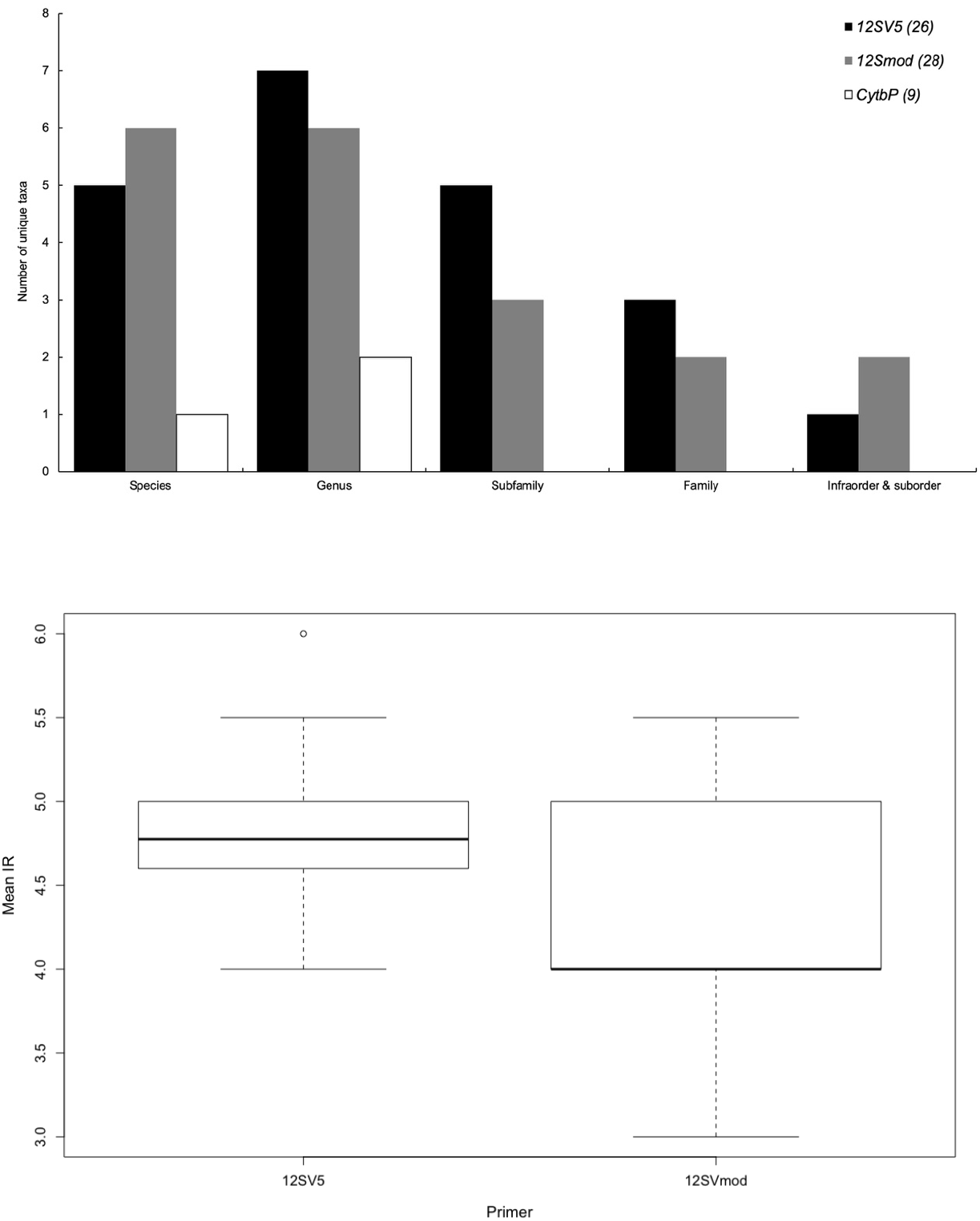
Primer-wise (top) taxonomic levels of the diet items identified using the three diet metabarcoding primers used in this study. The number samples which produced identifiable diet-specific sequences are indicated in parentheses next to each primer name; (bottom) The distribution of the mean taxonomic identification resolution (IR) among the samples represented by box plots, specific to the two better performing primers.

### 3.4 Diet composition

The number of diet-specific sequence variants in individual vulture faecal samples varied between 1 and 8 (2.73±0.33) for *12SV5* and between 1-6 (2.39±0.22) for *12SVmod*. This corresponded to a minimum of one diet taxon for each primer, and a maximum of five distinct diet species identified in a sample (GPN32) while using *12SV5*, and up to three diet species that could be distinguished in at least six samples using *12SVmod* (Fig. 3b). Across all samples, domestic cow was the most frequently occurring diet item, followed by spotted deer *Axis axis*. Some unexpected taxa were also identified including a free-tailed bat (family Molossidae; species found: *Tadarida* sp.) and a bent-winged bat (*Miniopterus* sp.), which were identified from faecal samples collected from cliffs where Long-billed vultures were nesting. Overall, the read counts did not represent the diet DNA concentrations in the mock community sample. Across replicates, blackbuck sequences were over-represented and barking deer sequences were very low in number, and not even retrieved using *12SVmod* and *CytbP* (Fig. 3a).

## 4. Discussion

We describe a robust dietary metabarcoding workflow for simultaneously uncovering the species identity, sex and diet of *Gyps* vultures, which are obligate scavengers of vertebrate carcasses, from non-invasively collected faecal samples. We could establish species identity and sex for all wild collected samples, which were at various stages of desiccation and collected from multiple substrates (leaves, dry and wet soil, sand, rocks), and recover diet sequences from all samples but one sample.

### 4.1 Performance of metabarcoding primers

The taxonomic coverage, resolution and number of false negatives in metabarcoding studies is strongly determined by the primer sets used (Pompanon et al., 2012). Our three sets of primers showed differential performance. *CytbV1* showed higher taxonomic resolution, while just *VsexZW* was best for sexing all the samples. Specifically, *CytbV1* successfully identified all the *Gyps* species and Red-headed vulture (subfamily Aegypiinae) unambiguously, but not Egyptian vulture (subfamily Gypaetiinae). Our goal was to include a primer that serves as the metabarcode for identifying the large Old World vultures belonging to the subfamily Aegypiinae, most of which are threatened with extinction (Buechley and Şekercioğlu, 2016), and *CytbV1* satisfied this criteria. Additionally, our sampling strategy which focused on collecting samples directly under a roost or nest appears to be effective since all field-collected unknown samples were found to belong to vultures. However, we found sequences belonging to two distinct species in one sample, most likely an outcome of mixing of nearby faecal droppings while collecting the sample from a site, where birds from both species were perched on a roost tree. This suggests the need for caution in identifying distinct droppings in future research.

Among the diet primers, *12SV5* and *12SVmod*, when used together, detected diet items in 29/30 samples. These two primers, targeting the same locus, also showed complementarity in taxa identified, and helped maximize taxonomic coverage and reduce false negatives as seen before while using the ‘one-locus-several-primer’ approach (Corse et al., 2019, 2017; Gibson et al., 2014). We used a blocking oligo with *12SV5*, which did not completely eliminate but depressed the amplification of vulture sequences and thereby, substantially increased the depth for identifying the diet items.

For future analyses of vulture characterization (species and sex) and diet, we recommend using a multiplex of *CytbV1, VsexZW, 12SV5* (with *GypsB*) and *12SVmod*. Sequences for vulture species identification and sexing, particularly specific to *VsexZW*, were retrieved in much higher numbers than diet-specific sequences (Table 1), possibly due to higher copy numbers in the samples. Furthermore, we amplified the nuclear markers separately and added a higher volume in the final amplicon mix in this run, since nuclear DNA is expected to be present in lower quantity than mitochondrial DNA. However, in future runs, we recommend reducing the concentration of *CytbV1* and *VsexZW* primers in the PCR reaction when multiplexing them with diet primers, or using lower volume of the PCR product if amplifying separately.

### 4.2 Importance of HTS filtering

The filtering pipelines most often used to analyse HTS metabarcoding data, including *mothur* (Schloss et al., 2009) and QIIME (Caporaso et al., 2010), rely on clustering reads into MOTUs. This often results in the number of taxa being overestimated (Brown et al., 2015; Clare et al., 2016). Instead, we used the clustering-free pipeline based on read counts of each sequence variant in each replicate (De Barba et al.2014; Corse et al. 2017). Here, non-arbitrary *LFN* thresholds based on positive and negative controls and replicates were used to filter the HTS data and minimize false positives (Fig. 1). This is a more conservative approach so that all variants with frequencies below the *LFN* thresholds are discarded, which might not be desirable when analysing complex communities, such as those collected for biodiversity assessments (e.g. Elbrecth & Leese 2017), where these filtering thresholds may have to be adequately relaxed. However, in our case, these thresholds appear appropriate since the dietary complexity is not very high (with the maximum number of species being six) in these obligate scavengers, which are known to specialize on the carcasses of large-bodied animals, especially ungulates (König, 1983; Mundy, 1992).

Furthermore, applying these *LFN* thresholds allowed us to retain most of the reads, while discarding a large number of potentially artefactual variants (Table 2). As a test, this filtering pipeline retrieved all expected taxa from the two positive controls we analysed (sample from captive individual; mock community sample). These also eliminated all variants from the negative controls, including potential contamination (human sequences) while using *12SV5* (0.2%) and greatly reduced it with *12SVmod* (0.06% from 4%). We also tested for tag jumps explicitly by including a negative control (*T_tag_*), which did not contain any reads after filtering. Moreover, we did not see the unique taxa from the mock community (blackbuck, barking deer) sequences in any of the wild collected samples, suggesting that mistagging bias (if any) was not appreciable. Comparing between PCR replicates, (where we discarded replicates using RD and retained only those variants which were present in at least two replicates) further helped minimize false positives (De Barba et al. 2014; Ficetola et al. 2016). It is also advisable to similarly include multiple biological replicates (distinct fractions from the same biological sample) to investigate the possibility of against false negatives (Lanzén et al., 2016; Zhan et al., 2014). Because the faecal samples are small in our case, we were unable to conduct more than one DNA extraction.

Finally, read counts serve as a poor estimator of diet species abundance/biomass in metabarcoding studies owing to PCR biases (Elbretch & Leese, 2015), which represents a limitation as compared to traditional dietary analyses. Even our data clearly indicated this limitation, wherein diet species added in the mock community (at equal extracted DNA concentration) were represented by skewed read counts. Some alternatives, such as using the number of sequence variants for every taxon as an index of prey biomass (Jo et al. 2016) or minimum number of individuals consumed (Corse et al. 2017), have been suggested. However, for the mock communities, we retrieved multiple variants for every taxon although DNA from just one individual was added, making these approximations unreliable. More careful evaluations using appropriate cafeteria trials and mock samples as controls are required to establish how to convert read counts reliably into relative biomass consumed.

### 4.3 Identification of diet composition

The metabarcodes used in our study provided a high level of taxonomic resolution and we could identify most diet items to the species level, particularly when supplemented with BLAST hits and biogeographic information on plausible occurrence in the study sites. However, we were unable to confirm the species-level identity of the bat sequences recovered owing to both paucity of complete species inventory from the study localities and lack of representation in reference databases. Overall, the metabarcodes appear to be adequate for the overall goal of our study, which is to be able to differentiate between large domestic livestock, which receive veterinary care and pose the threat of diclofenac, from wild ungulates and small domesticated animals.

From the small sample of wild collected faecal samples, it appears that domestic cow (and buffalo) are the dominant dietary item, in sites both inside and outside National Parks (Fig. 3b). This may be attributed to the generally high availability of cow carcasses in India, where some of the largest numbers of livestock are found and cows are not slaughtered owing to cultural taboo (Cuthbert et al. 2016). However, samples from sites inside National Parks also contained wild species including spotted deer. Other diet species included other ungulates including Dromedary Camel, Nilgai, primate (Rhesus macaque), a *Canis sp*. and Indian peafowl. We also recovered bat sequences from some faecal samples collected from Indian vultures, which nest on steep cliffs where bats often roost. While they could represent environmental contamination, given that these were not present in all the samples collected from the same sites and were represented by high number of reads, we presume that is unlikely. However, to conclusively comment on whether it was really ingested or is an environmental contaminant, we suggest collecting field negative controls in the future (Zinger et al. 2019). Finally, establishing whether access to carcasses of wild animals from National Parks is really impacting the risk of diclofenac-poisoning will require analysing many more samples representing both spatial and temporal spread of the dietary ecology of these species, while accounting for other ecological factors such as habitat and the relative availability of wild ungulates and domestic livestock.

## 5. Conclusion

Dietary analyses generate critical information regarding trophic interactions and food/habitat relationships. This is particularly important for the Critically Endangered *Gyps* species in the Indian subcontinent, where such information is required for assessing the risk of diclofenac poisoning and identifying potentially safe refuges where captive stock can be released for best conservation outcomes. Veterinary diclofenac was banned in 2006. However, diclofenac formulated for human use continues to be used illegally and several other veterinary drugs (e.g. ketoprofen, aceclofenac, nimesulide) available for legal use are known to be nephrotoxic to be vultures (Cuthbert et al., 2016; Galligan et al., 2016; Naidoo et al., 2010). Hence, understanding the contribution of domestic livestock in the diet of vultures could still serve as an important index of mortality risk.

Our results establish that the robust DNA metabarcoding workflow described here can be efficiently utilised to describe the species- and sex-specific diet of wild vultures from non-invasively collected faecal samples, collected across a large geographic scale. We followed an experimental design that included multiple controls and allowed us to filter the HTS data using non-arbitrary thresholds to minimize false-positives. The stringent filtering pipeline yielded a robust data set that guaranteed high confidence in diet species identification and enabled vulture and diet characterization in almost all samples. Finally, we hope to use this “field-to-benchtop-to-desktop” workflow to generate robust rangewide dietary data to facilitate evidence-based conservation decision-making to recover populations of these imperilled obligate scavengers.

## Acknowledgements

We wish to thank the state forest departments of Madhya Pradesh, Haryana and Rajasthan for providing research permissions for collecting samples. The Raptor Research and Conservation Fund (RRCF-23/08/19) is thanked for funding. MGH is funded through a NCBS-inStem-Cambridge postdoctoral fellowship. William Amos is thanked for useful inputs and discussion on the drafts. We thank Himanshu Chhattani and Kaushal Patel for help in field. Meghana Natesh is thanked for useful discussions on laboratory protocol. Anuradha Savand is thanked for help with lab work. The Next-Generation-Sequencing facility at NCBS is thanked for processing the samples.

## Author Contributions section

MGH, RG and UR conceived and designed the study. VP provided samples from captive vultures and provided key insights on vulture ecology. MGH and NG collected the samples from the wild. MGH, NG, IS and AP performed the laboratory experiments. MGH and HS analysed the data. MGH wrote the manuscript and all authors helped to draft and improve the manuscript. All authors read and approved the manuscript.

