## Supplementary file for "Metabarcoding for parallel identification of species, sex and diet: an application to the conservation of globally-threatened *Gyps* vultures"

**Supplementary material**

**Table S1. Potential species that *Gyps* vultures scavenge on across Indian subcontinent.**

| English name | Scientific name |
|---|---|
| **Bovidae** | |
| Banteng | *Bos javanicus* |
| Indian cow | *Bos indicus* |
| European cow | *Bos taurus* |
| Gaur | *Bos gaurus* |
| Yak | *Bos grunniens* |
| Domestic water buffalo | *Bubalus bubalis* |
| Wild water buffalo | *Bubalus arnee* |
| Goat | *Capra hircus* |
| Markhor | *Capra falconeri* |
| Sheep | *Ovis aries* |
| Blue sheep | *Pseudois nayaur* |
| Himalayan tahr | *Hemitragus jemlahicus* |
| Nilgiri tahr | *Nilgiritragus hylocrius* |
| Ghoral | *Naemorhedus goral* |
| Takin | *Budorcas taxicolor* |
| Chinkara | *Gazella bennetii* |
| Blackbuck | *Antilope cervicapra* |
| Nilgai | *Boselaphus tragocamelus* |
| Four-horned antelope | *Tetracerus quadricornis* |
| **Cervidae** | |
| Spotted deer | *Axis axis* |
| Hog deer | *Hyelaphus porcinus* |
| Red deer | *Cervus elaphus* |
| Barasingha | *Rucervus duvaucelii* |
| Sambar | *Rusa unicolor* |
| Muntjac | *Muntiacus muntjak* |
| Manipur brow-antlered deer | *Rucervus eldii* |
| Chevrotain | *Moschiola indica* |
| **Camelids** | |
| Dromedary | *Camelus dromedarius* |
| Bactrian camel | *Camelus bactrianus* |
| **Suidae** | |
| Wild boar | *Sus scrofa cristatus* |

| | |
|---|---|
| Domestic pig | *Sus scrofa domesticus* |
| **Perissodactyla** | |
| Horse | *Equus caballus* |
| Donkey | *Equus asinus* |
| Wild ass | *Equus hemonius* |
| Greater one-horned rhinoceros | *Rhinoceros unicornis* |
| Indian Elephant | *Elephas maximus* |
| **Carnivora** | |
| Dhole | *Cuon alpinus* |
| Golden jackal | *Canis aureus* |
| Dog | *Canis familiaris* |
| Wolf | *Canis lupus* |
| Leopard | *Panthera pardus* |
| Tiger | *Panthera tigris* |
| Sloth bear | *Melursus ursinus* |
| Himalayan bear | *Ursus thibetanus* |
| Sun bear | *Helarctos malayanus* |
| **Primates** | |
| Macaque | *Macaca sp.* |
| Langur | *Semnopithecus sp.* |
| Rodents | |
| Marmot | *Marmota sp.* |

**Figure S1.** Metabarcode region and primer positions (12S mitochondrial gene) for the potential diet species (from NCBI nucleotide database).

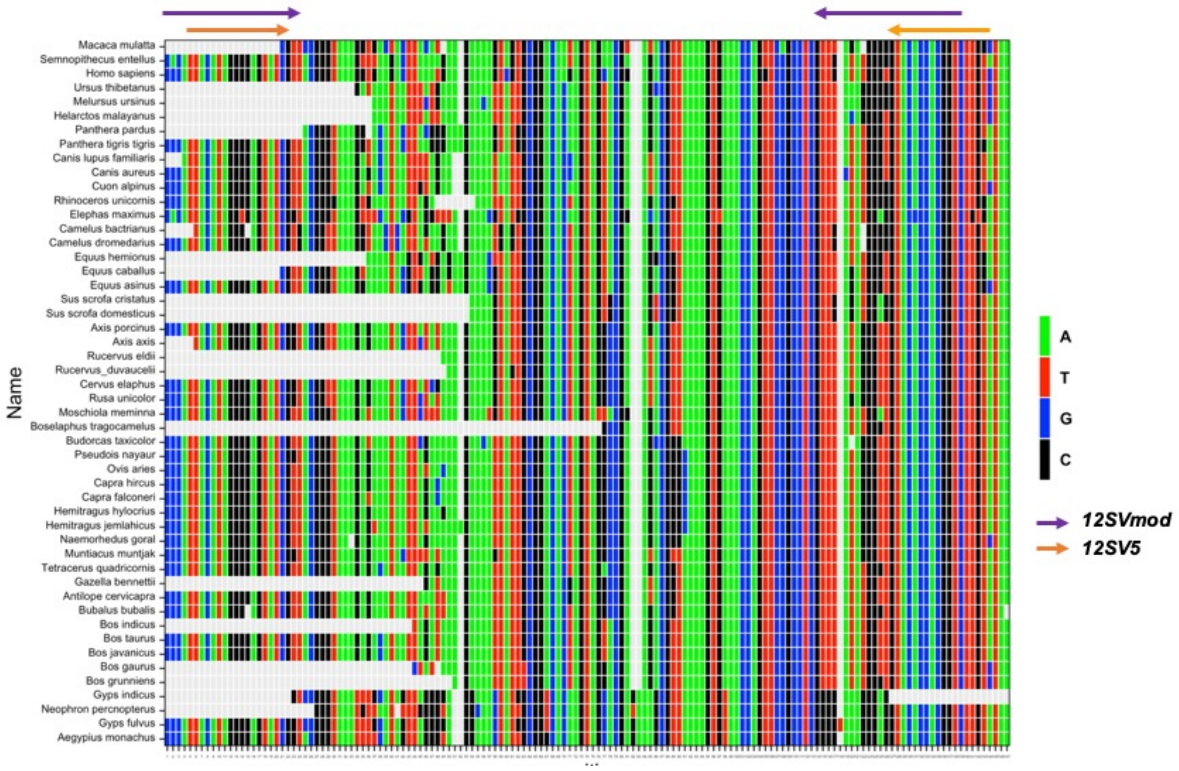

**Figure S2.** Schematic representation of the benchtop workflow for DNA metabarcoding.

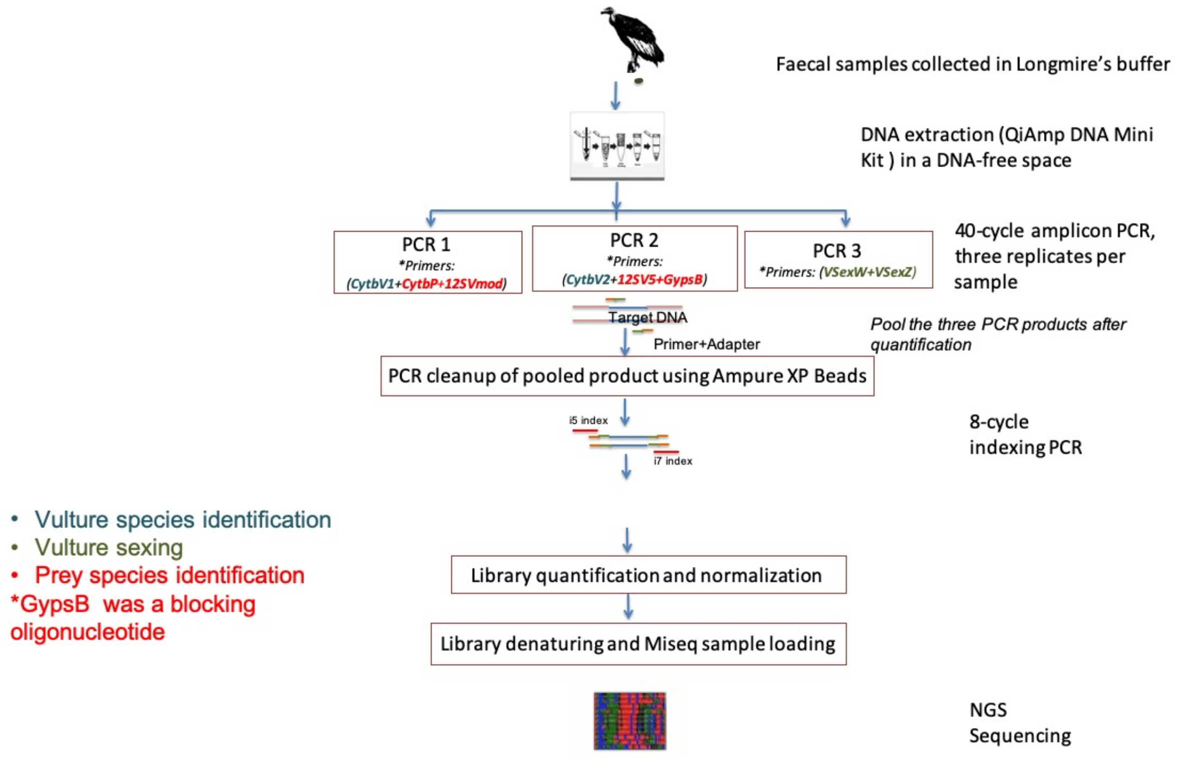
